# A Conserved Regenerative Architecture Underlies Skeletal Muscle Repair in Adult Zebrafish

**DOI:** 10.64898/2026.08.12.744335

**Authors:** Mirjana Novkovic, Andjela Milicevic, Emilija Milosevic, Luka Bojic, Jovana Jasnic, Snezana Kojic

## Abstract

Adult zebrafish efficiently regenerate skeletal muscle following different types of injury; however, the molecular programs involved in repair after extensive cryoinjury remain to be comprehensively characterized. Here, we explored the transcriptomic response of adult zebrafish skeletal muscle at 7 days post cryoinjury (dpci), a stage marked by ongoing tissue clearance, progenitor expansion, myogenic differentiation, and nascent myofiber formation, and compared it with phase-matched stab wound injury. Cryoinjury induced a broad transcriptional response, with 5,330 differentially expressed genes. Integrated enrichment and protein–protein interaction analyses revealed that, at 7 dpci, zebrafish skeletal muscle functions as an integrated regenerative system in which immune remodeling, progenitor expansion, myogenic differentiation, extracellular matrix reconstruction, mechanotransduction, biosynthetic adaptation, proteostasis, and intracellular trafficking operate simultaneously. In parallel, mature sarcomeric and oxidative metabolic programs were suppressed, consistent with ongoing tissue reconstruction and structural immaturity. Comparison with stab-wounded skeletal muscle revealed substantial transcriptional conservation, as 612 of 717 stab-wound-responsive genes (85%) were also differentially expressed after cryoinjury. Shared upregulated genes formed coherent functional modules related to proliferation, extracellular matrix organization and signaling, immune regulation, muscle differentiation, and protein processing. Thus, distinct injury modalities converge on a common regenerative program, while cryoinjury elicits a quantitatively broader transcriptional response. These findings support a conserved regenerative architecture of adult zebrafish skeletal muscle repair, in which interconnected biological modules act coordinately, with the breadth of transcriptional engagement reflecting regenerative demand.

## INTRODUCTION

Skeletal muscle repair is a highly coordinated biological process in which damaged tissue is progressively cleared, rebuilt, and functionally restored. Following injury, skeletal muscle healing is commonly described as a sequence of partially overlapping phases of degeneration, inflammation, regeneration, and remodeling (Laumonier and Ménétrey 2016; Endo 2023) requiring tightly coordinated interactions among multiple cell types and extracellular matrix (ECM) components (Hardy et al. 2016). The extent and quality of skeletal muscle repair depend strongly on the type, magnitude, and microenvironment of injury. These differences are particularly important in traumatic injuries, where tissue architecture can be profoundly disrupted, and functional recovery may remain inadequate (Warren et al. 2007; Hardy et al. 2016).

A broad range of experimental injury models has been developed to study skeletal muscle repair, including myotoxin-induced, chemical, mechanical, and physical injuries (Warren et al. 2007; Hardy et al. 2016). Although these models involve the same general phases of regeneration, they differ in the extent of muscle fiber damage, preservation of the tissue environment, inflammatory response, ECM remodeling, and final repair outcome (Hardy et al. 2016). Freeze injury is a useful model because it produces a clearly defined area of necrotic damage and reproducible tissue disruption, allowing degeneration and repair to be studied in a controlled way (Le et al. 2016). In contrast, volumetric muscle loss represents a distinct condition characterized by irreversible tissue loss, persistent fibrosis, and chronic functional deficits (Nuutila et al. 2017; Whitaker et al. 2025). Comparative studies in mammals have shown that different injury paradigms trigger both shared and injury-specific transcriptional programs and have a major influence on the subsequent regenerative response (Hardy et al. 2016). In mammalian models, gene-expression studies have revealed that freeze injury is associated with strong inflammatory, angiogenic, and tissue-remodeling signatures (Warren et al. 2007), whereas non-regenerative injuries such as volumetric muscle loss show prolonged inflammation, pro-fibrotic signaling, and impaired activation of pro-myogenic pathways (Whitaker et al. 2025).

Although mammalian models remain indispensable, zebrafish have emerged as a powerful complementary system for skeletal muscle research (Lu et al. 2025). One of the major advantages of zebrafish is their ability to maintain lifelong muscle growth through continued recruitment of new myofibers (Lu et al. 2025). Spatial transcriptomic analysis of adult zebrafish muscle has further shown that musculature is regionally heterogeneous and retains transcriptional signatures consistent with ongoing myogenic activity (Liu et al. 2022). These features make zebrafish especially valuable for studying vertebrate skeletal muscle response to injury and regulation of adult muscle growth, regeneration, and stem-cell dynamics.

Studies in zebrafish have already demonstrated that muscle regeneration involves Pax7-positive satellite-like cells and other myogenic progenitors, supporting the view that adult zebrafish muscle regeneration relies on conserved stem-cell-mediated mechanisms (Knappe et al. 2015; Berberoglu et al. 2017). In addition, the regenerative response is influenced by developmental stage and injury extent (Knappe et al. 2015). Minor injuries such as stab wounds or laser-induced damage are followed by efficient repair in larvae and adults (Otten and Abdelilah-Seyfried 2013; Knappe et al. 2015; Berberoglu et al. 2017). However, until recently, it remained unclear whether adult zebrafish could restore profoundly damaged locomotory musculature after extensive injury. This gap has now been addressed by the development of a cryoinjury model targeting the caudal peduncle, in which several myomeres undergo severe localized destruction while preserving body integrity (Oudhoff et al. 2024). Using this model, Oudhoff and colleagues showed that adult zebrafish can regenerate extensively cryoinjured skeletal muscle within approximately one month, with wound clearance, transient connective tissue deposition, proliferation of muscle stem cells, myogenic differentiation, and progressive restoration of slow and fast muscle compartments (Oudhoff et al. 2024). However, the regenerated tissue does not fully recapitulate the original pattern, indicating that even in this highly regenerative vertebrate, tissue restoration remains incomplete (Oudhoff et al. 2024).

Despite these advances, the global transcriptional response of adult zebrafish skeletal muscle to cryoinjury remains uncharacterized. Histological and cellular analyses have established that this model supports robust regeneration after severe injury (Oudhoff et al. 2024), but a transcriptome-level view is needed to define the molecular programs that accompany tissue healing. Such analysis is important not only for understanding zebrafish muscle regeneration itself, but also for positioning this model within the broader landscape of vertebrate muscle injury systems (Warren et al. 2007; Lu et al. 2025).

The present study aimed to investigate the transcriptomic response of cryoinjured skeletal muscle in adult zebrafish at 7 days post-cryoinjury (dpci), corresponding to a critical stage of repair when newly formed muscle fibers begin to infiltrate the nascent connective tissue at the injury site, replacing damaged fibers (Oudhoff et al. 2024). Comparison with phase-matched stab-wounded skeletal muscle was designed to determine whether distinct injury paradigms converge on common regenerative programs.

## METHODS

### Zebrafish husbandry

Zebrafish (AB strain) were maintained in a circulating water system, at 28°C, under a 14h light:10h dark cycle, in accordance with institutional and national ethical and animal welfare guidelines, harmonized with EU Directive 2010/63/EU (amended by Directive 2024/1262). Experiments were approved by the Veterinary Directorate, Ministry of Agriculture, Forestry and Water Management, Republic of Serbia (No. 000094465 2024).

### Skeletal muscle injury and sample preparation

Cryoinjury of the zebrafish skeletal muscle located in the caudal region was performed following the procedure reported by Oudhoff et al (Oudhoff et al. 2023). In brief, zebrafish were anesthetized in system water containing 0.6 mM buffered tricaine (MS-222 ethyl-m-aminobenzoate, Sigma-Aldrich). Skeletal muscle cryoinjury was then induced using a stainless-steel cryoprobe, prepared according to the established protocol. The cryoprobe was precooled in liquid nitrogen and applied for 6 seconds to the skin on the left side of the fish, positioned perpendicular to the body surface between the anal and caudal fins. Following injury, fish were immediately returned to system water and monitored continuously until normal swimming behavior resumed. Following the recovery period, tissue samples from the injured caudal peduncle were harvested at 7 dpci. Stab injury was performed as previously described (Milovanovic et al. 2025).

### Isolation of RNA

Skeletal muscle tissue around the cryoinjury (1-2 mm from the injury edges), stab wound or from control non-injured zebrafish was dissected out, deskinned, and deboned. Tissue was homogenized and processed in TRI Reagent (Invitrogen, Thermo Fisher Scientific), and total RNA was extracted using RNA Clean and Concentrator Kit (ZYMO Research), all following the manufacturer’s instructions. RNA purity and concentration were assessed using BioSpec-nano (Shimadzu) by measuring absorbance at 260 and 280 nm and Qubit 4^TM^ Fluorometer with ssRNA BR Assay Kit^TM^ (Thermo Fisher Scientific). RNA samples were stored at -80°C and used for downstream analysis.

### Reverse transcription quantitative polymerase chain reaction (RT-qPCR)

Complementary DNA (cDNA) was synthesized from total RNA using the High-Capacity cDNA Reverse Transcription Kit (Applied Biosystems), according to the manufacturer’s protocol, with 1 µg of RNA as a template. Amplification was carried out using a Power SYBR™ Green PCR Master Mix (Applied Biosystems) and Line-Gene 9600 Plus real-time PCR system (Bioer Technology) under the following conditions: initial denaturation at 95°C for 10 min, followed by 40 cycles of 95°C for 15 s and 60°C for 1 min. Melting curve analysis was performed to confirm amplification specificity. Data were analyzed by the −ΔΔCt method. Reactions were performed in technical triplicate for each sample. The Ct value of each gene was normalized to the Ct value of *rpl13a,* which was used as an internal control. The primer sequences are listed in Supplementary Table S1. Gene expression levels in each experimental group were normalized to those of the non-injured control group. Statistical analysis was conducted using GraphPad Prism software 6.0. For comparisons between injured and non-injured groups, the unpaired two-tailed Student’s t-test was used.

### RNA sequencing

Bulk RNA-seq was performed using RNA isolated from cryoinjured skeletal muscle at 7 dpci (n=4). The quality of RNA was determined using the RNA 6000 Nano Kit (Agilent Technologies) and Bioanalyzer 2100 (Agilent Technologies). All samples had an RNA integrity number (RIN)≥ 5.5, concentration ≥ 50 ng/µl, and OD 260/280 between 1.8 and 2.1, and were sequenced at Novogene (Munich, Germany).

The cDNA libraries were constructed via poly A enrichment using the Illumina TruSeq Stranded mRNA Library Preparation kit (Illumina), and 150 paired-end sequencing was performed on the SEQ Novaseq 6000 platform (Illumina).

To assess transcriptomic changes following skeletal muscle cryoinjury, previously sequenced RNA samples from non-injured skeletal muscle (dataset GSE277480) were used as controls. To minimize processing bias, FASTQ files from both control and cryoinjured samples were analyzed together. Quality assessment of raw reads was performed using FastQC v0.12.1, and read trimming and filtering were conducted with fastp v0.23.4 to remove adapter sequences and retain high-quality reads (parameters: qualified_quality_phred=20, nqualified_percent_limit=30, average_qual=25, low_complexity_filter=True, complexity_threshold=30). Clean paired-end reads were aligned to the zebrafish reference genome (GRCz11, Ensembl v112) using STAR v2.7.11b, with the sjdbOverhang parameter set to 149 bp and all other parameters left as default. The percentage of uniquely mapped reads was high, with an average of 85.4% (range: 83.7 - 87.4%). Gene-level abundance estimates were generated using RSEM v1.3.3, and sample-specific results were combined into a single raw expression matrix using the rsem-generate-data-matrix command.

The RNA-seq data from cryoinjured skeletal muscle are deposited in NCBÍs Gene Expression Omnibus and are accessible through the GEO Series accession number GSE319584, available at https://www.ncbi.nlm.nih.gov/geo/query/acc.cgi?acc=GSE319584. RNA-seq data sets of non-injured and stab-wounded skeletal muscle at 5 days post injury (dpi) are available under accession number GSE277480.

### Differential gene expression and enrichment analysis

The raw expression matrix was normalized and filtered to include only genes with at least 10 counts in n samples, where n corresponds to the sample number in the smallest experimental group (n=4). Normalization and filtering were performed using the edgeR (v4.4.1) program package in the R environment (v4.4.2).

Exploratory data analysis was conducted using a box plot and principal component analysis (PCA). To correct for unwanted variation arising from differences in library preparation time and sequencing facilities (despite identical Illumina technology and kits), the RUVg function from the RUVSeq v1.40.0 package was used. Negative control genes were defined as those with less than 20% variance (standard deviation/mean) across all samples and mean expression greater than 5 counts. One factor of unwanted variation was estimated and incorporated into the model for the analysis of differentially expressed genes (DEGs).

Differential expression analysis between cryoinjured and non-injured groups (2 groups, 4 samples each) was performed using a multifactorial negative binomial generalized linear model computed by edgeR (v4.4.2). P-values were adjusted for multiple testing using the Benjamini & Hochberg method, and genes with an adjusted p≤ 0.05 and an absolute log_2_ fold change≥ 1 were considered significantly differentially expressed. The bioMart package v.2.62.0 was used to annotate DEGs, querying available Ensembl Gene IDs and retrieving Entrez gene IDs, Uniprot accessions, and ZFIN gene symbols.

Gene Ontology (GO) annotation and Kyoto Encyclopedia of Genes and Genomes (KEGG) pathway annotation of DEGs were performed separately for up- and down-regulated genes using clusterProfiler v4.14.4 and org.Dr.eg.db as a reference database. Gene set pathway enrichment analysis was conducted using ReactomePA v1.50.0. Enriched terms and pathways with an adjusted p≤ 0.05 were considered significantly enriched. The results were visualized using R packages gplots v3.2.0 and ggplot2 v3.5.1. A list of 15106 genes, remaining after normalization and filtering of raw counts, was used as the reference background set for all enrichment analyses.

### Protein-protein interaction network analysis

DEGs were converted into a STRING protein–protein interaction (PPI) network (Szklarczyk et al. 2022). The network was analyzed in Cytoscape v3.10.4 (Shannon et al. 2003) using the MCODE plugin v2.0.3 with default parameters. Based on the MCODE score, up to five gene clusters were selected for functional enrichment analysis against Gene Ontology (Biological Process, Molecular Function, and Cellular Component), KEGG, Reactome, and WikiPathways databases, using a similarity cutoff of 0.7. To facilitate biological interpretation and reduce redundancy among overlapping enrichment terms identified across multiple annotation databases, significantly enriched terms were manually consolidated into broader functional (umbrella) terms (Supplementary Tables S6 and S10). Terms describing complementary aspects of the same biological process, molecular function, cellular component, or signaling pathway were grouped according to their shared biological context and functional relationships, irrespective of their source database. These umbrella terms served exclusively as an interpretative framework for describing the major biological themes represented within individual PPI clusters and did not influence the underlying enrichment analyses. The results were visualized using ggplot2 v3.5.1 and Cytoscape v3.10.4.

## RESULTS

### Cryoinjury causes an extensive transcriptomic response in skeletal muscle

To identify genes and pathways involved in the repair of adult zebrafish skeletal muscle, transcriptome profiling was performed at 7 dpci. Non-injured skeletal muscle was used as a control. PCA showed that cryoinjury significantly impacts gene expression profile, with clear separation between injured and non-injured samples (PC1=48,4%; PC2=12% of total variance), reflecting relevant biological differences (Fig. 1A).

**Fig. 1.**
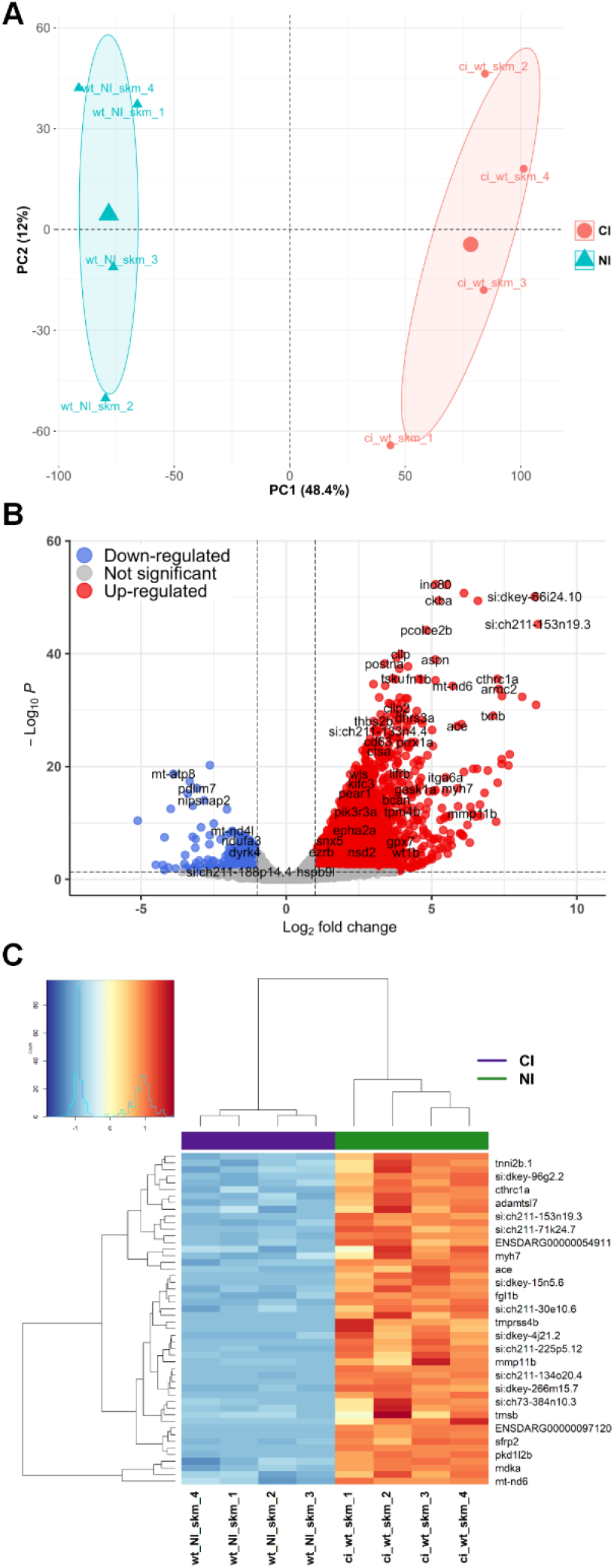
Transcriptomics analysis of cryoinjured zebrafish skeletal muscle at 7 dpci. (**A**) PCA plot depicting transcriptome heterogeneity between samples (n=4 for each experimental group); (**B**) Volcano plot indicating DEGs between cryoinjured and non-injured adult skeletal muscle; red dots, upregulated in cryoinjury; blue dots, downregulated in cryoinjury; gray dots, nonsignificant genes. (**C**) Heatmap of the top 50 DEGs between cryoinjured and non-injured adult skeletal muscle showing expression patterns across samples. The Z-scores of gene expression measurements are displayed as colors ranging from blue to red. NI, non-injured; CI, cryoinjured.

A total of 5330 genes were found to be significantly differentially expressed at least 2 times (|log2 fold change|≥1), indicating that these genes might be associated with the regulation of skeletal muscle repair after cryoinjury. Cryoinjury caused upregulation of 5077 and downregulation of 253 genes (Supplementary Table S2). Among upregulated DEGs, several myogenic regulatory factors have been found, including *myf5* and *myog*, as well as embryonic myosin *myh7*, while myozenin (*myoz1a/b* and *myoz2a/b*) and troponin (*tnnt2c*, *tnnt3a*) genes were significantly downregulated. The volcano plot of all genes and the heatmap of the 50 most variable genes are shown in Fig. 1B and C.

To elucidate the molecular function and biological processes in which DEGs are involved, we performed Gene Ontology (GO) overrepresentation analysis. GO enrichment was conducted separately for upregulated and downregulated genes (Supplementary Table S3). The analysis revealed that upregulated genes were significantly associated with cell adhesion and migration, mitotic cell division, and cartilage, connective tissue, and skeletal system development (Fig. 2A, Supplementary Table S3). Enrichment of GO terms related to tissue regeneration and developmental growth morphogenesis was also observed among the upregulated genes (Fig. 2A, Supplementary Table S3). In contrast, regulation of metabolites and energy processes, as well as contractile muscle fiber and sarcomere organization (Fig. 2B, Supplementary Table S3), were enriched among downregulated genes.

**Fig. 2.**
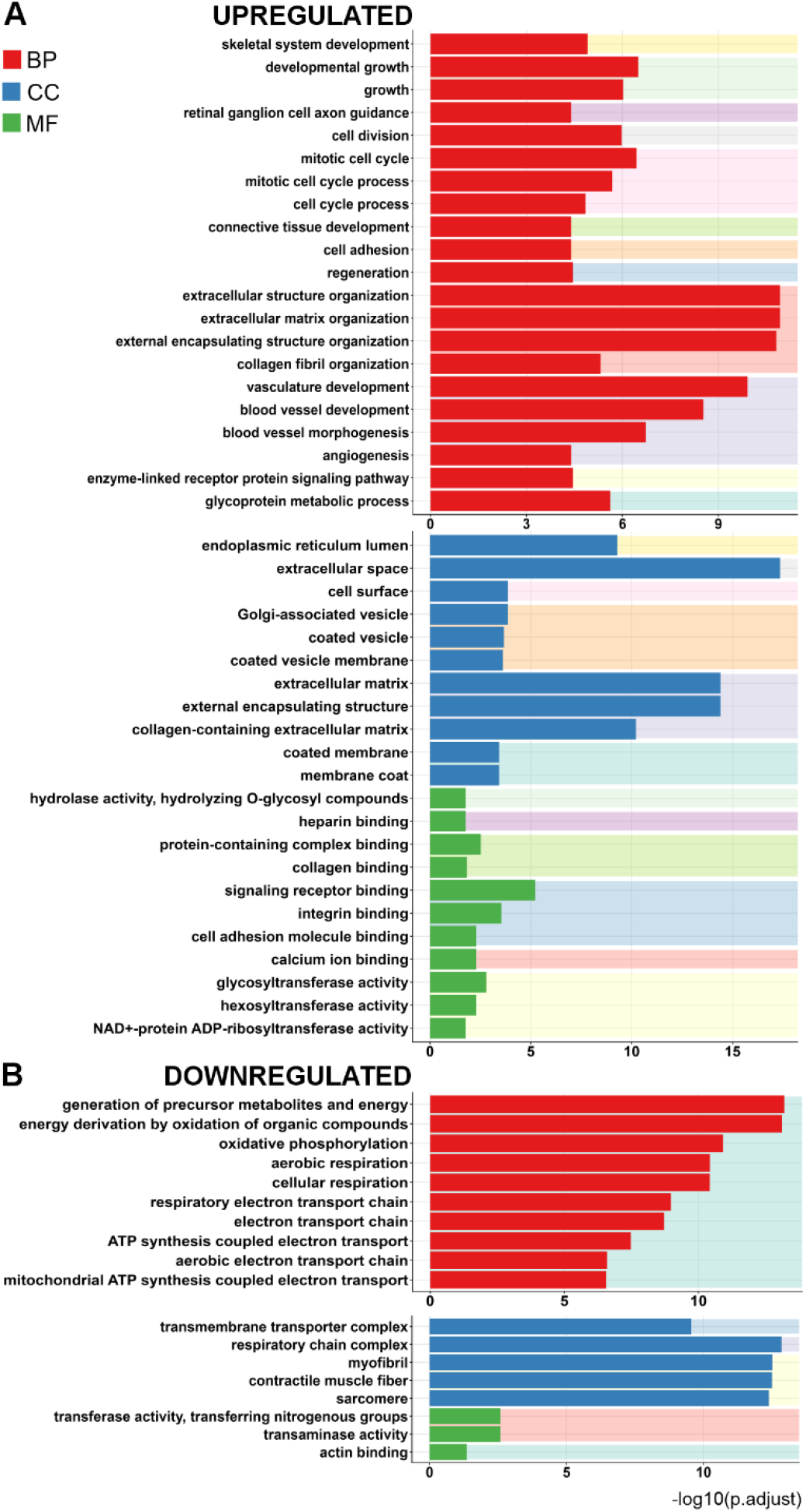
Gene ontology analysis of upregulated (**A**) and downregulated (**B**) genes in biological processes (BP, red bars), cellular components (CC, blue bars), and molecular function (MF, green bars). Bars with the same background color denote clusters of related GO terms.

### Pathway and network analyses point to broad functional reorganization of cryoinjured skeletal muscle at 7 dpci

To gain insight into the biological pathways associated with the transcriptional response to cryoinjury at 7 dpci, we performed KEGG (Kyoto Encyclopedia of Genes and Genomes) and Reactome pathway enrichment analyses of DEGs. KEGG pathway analysis revealed significant enrichment of upregulated genes in pathways related to regulation of actin cytoskeleton, focal adhesion, ECM-receptor interaction, apoptosis, and TGFβ signaling, as well as to lysosome and phagosome (Fig. 3A, Supplementary Table S4). Downregulated genes were significantly enriched in multiple metabolic pathways, including oxidative phosphorylation, glycolysis/gluconeogenesis, and starch and sucrose metabolism, together with pathways specific for mature muscle fibers such as cardiac muscle contraction and cytoskeleton in muscle cells (Fig. 3A, Supplementary Table S4), consistent with ongoing myocyte differentiation during muscle recovery (Montfort et al. 2016). These findings were complemented with Reactome pathway analysis which showed enrichment of similar pathways among upregulated genes, including innate immune and inflammatory response pathways (innate immune system, RHO GTPase effectors, neutrophil degranulation, RHO GTPases activate formins, MHC class II antigen presentation, platelet degranulation), collagen extracellular biosynthesis, axon biology signaling development and regulation of actin dynamics related to phagocytosis (Fig. 3B, Supplementary Table S5). Downregulated genes contributed to enrichment of aerobic respiration and respiratory electron transport (Supplementary Table S5).

**Fig. 3.**
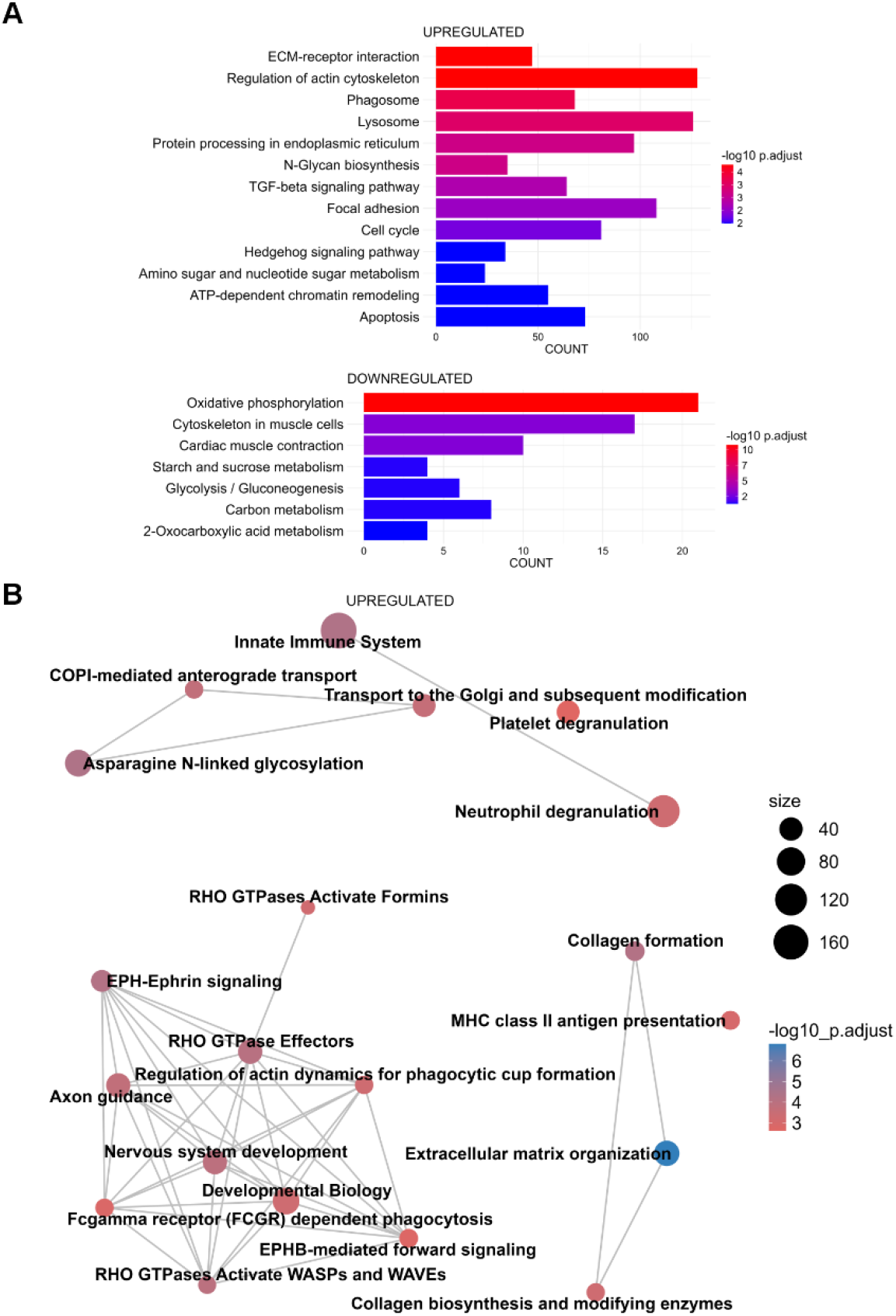
Overrepresentation analysis of DEGs in cryoinjured skeletal muscle: (A) KEGG pathway analysis of up- and downregulated genes; (B) Reactome pathway analysis of upregulated genes. Size - number of genes significantly enriched in the pathway.

PPI network analysis of DEGs upregulated at 7 dpci in cryoinjured adult zebrafish skeletal muscle identified five major functional clusters (Fig. 4 and Supplementary Table S6). Cluster 1 (score 50.82; 55 genes) was enriched for ribosome biogenesis and RNA processing, including rRNA maturation, snoRNA-guided modification, and assembly of preribosomal complexes within the nucleolus. Cluster 2 (score 27.43; 50 genes) captured mitotic cell cycle progression, encompassing DNA replication, spindle organization, and chromosome segregation. Cluster 3 (score 22.22; 126 genes) integrated cell cycle progression, RNA processing and ECM interaction, including spliceosomal activity, DNA replication machinery, and collagen/integrin-associated components. Cluster 4 (score 16.87; 185 genes) was enriched for cytoskeletal remodeling and intracellular transport, including Rho GTPase signaling, actin dynamics, vesicle trafficking, nucleocytoplasmic transport, and proteasome-associated processes. Cluster 5 (score 11.64; 134 genes) captured signal transduction pathways and membrane-associated processes, including phosphatidylinositol 3-kinase (PI3K) complexes, mTOR, Hedgehog, and insulin signaling, and vesicular transport. Downregulated DEGs formed four major functional clusters (Supplementary Fig. S1 and Supplementary Table S6), mainly reflecting suppression of metabolic programs. Cluster 1 (score 17.56; 19 genes) was strongly enriched for mitochondrial oxidative metabolism, including oxidative phosphorylation, respiratory chain complexes, and ATP synthase activity. Cluster 2 (score 3.33; 4 genes) highlighted endoplasmic reticulum (ER)–associated processes, including regulation of nitric oxide synthase activity. Cluster 3 (score 3.00; 7 genes) encompassed central carbon and intermediary metabolism, including glycolysis/gluconeogenesis, amino acid biosynthesis, and 2-oxocarboxylic acid metabolism. Cluster 4 (score 3.00; 3 genes), associated with myosin filament organization, points to reduced expression of a few genes encoding components of the contractile apparatus.

**Fig. 4.**
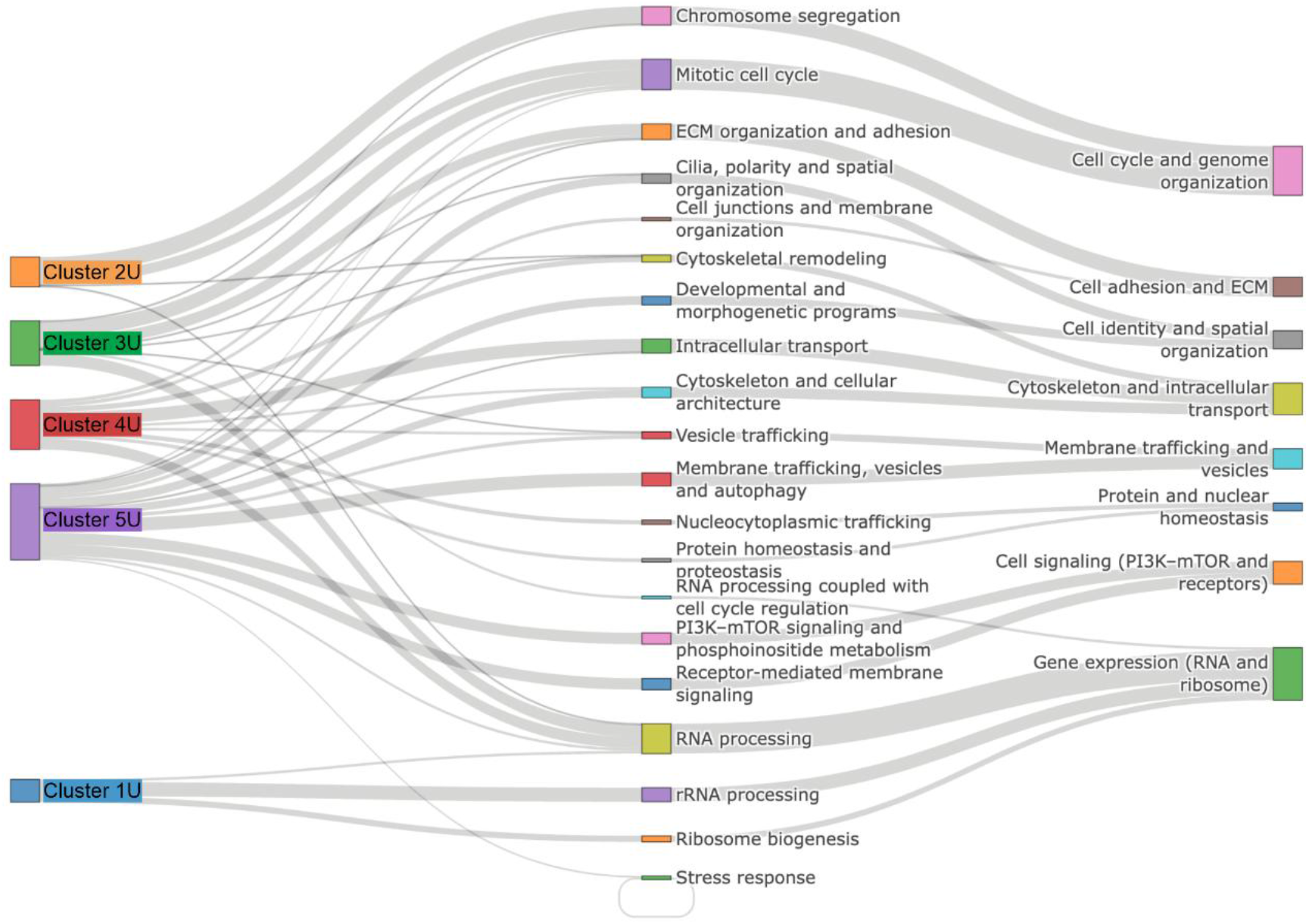
Functional relationships of upregulated DEGs in cryoinjured adult zebrafish skeletal muscle at 7 dpci. Sankey diagram showing clusters derived from PPI network analysis of DEGs and their associated GO, KEGG, Reactome or Wiki Pathways enrichment terms. Flow widths represent the number of genes linking each cluster to enriched terms. Cluster 1U: Nucleolar ribosome biogenesis and rRNA processing; Cluster 2U: Cell proliferation and mitotic machinery; Cluster 3U: Cell cycle–coupled RNA processing and ECM remodeling; Cluster 4U: Intracellular organization: cytoskeleton, trafficking, and proteostasis, and Cluster 5U: PI3K–mTOR signaling, membrane trafficking, and developmental morphogenesis.

### Comparative transcriptomic analysis reveals a shared regenerative program in cryoinjured and stab-wounded zebrafish skeletal muscle

To investigate transcriptional response of skeletal muscle to different types of injury, we compared RNA-seq profiles of cryoinjured (this manuscript) and stab-wounded (GSE277480) skeletal muscle at comparable regenerative stages (7 dpci for cryoinjury and 5 dpi for stab wound). According to RNA-seq results (Fig. 5A), and confirmed by RT-qPCR (Fig. 5B), *myog* (myogenin, myogenic regulatory factor that promotes terminal differentiation of myoblasts), *myh7* (embryonic/developmental myosin heavy chain isoform, a marker of newly formed myofibers), and *mymk* (myomaker, factor essential for fusion of differentiated myoblasts into multinucleated myotubes) were upregulated, indicating progression of skeletal muscle regenerative program into the myogenic differentiation, myoblast fusion, and formation of nascent myofibers. At the histological level, newly forming myofibers were visible in wounded tissues at designated time points (Fig. 3D-F’’ in Milovanovic et al. (Milovanovic et al. 2025) and Fig. 3E in Oudhoff et al. (Oudhoff et al. 2024)).

**Fig. 5.**
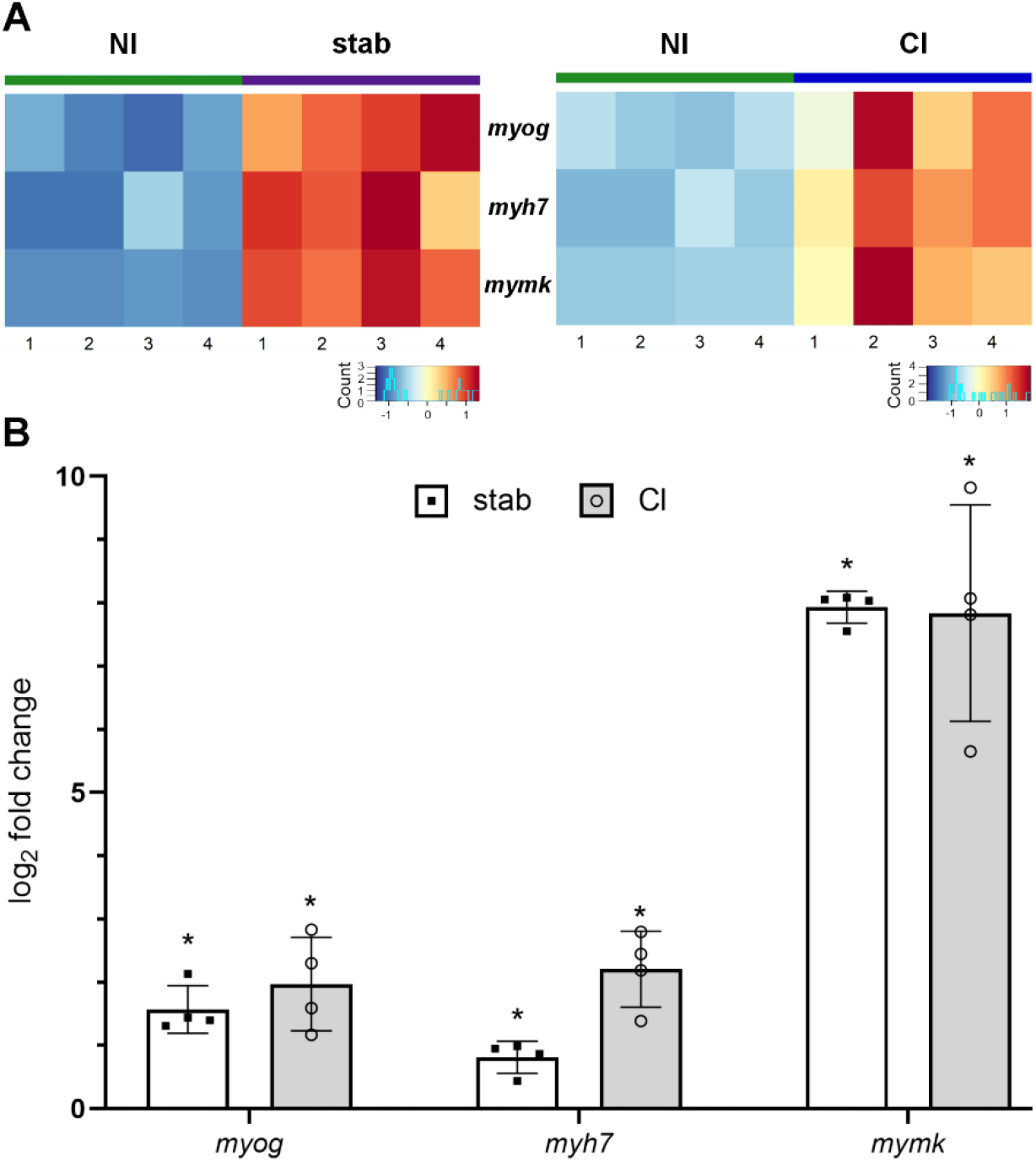
Stab-wounded (at 5 dpi) and cryoinjured (at 7 dpci) skeletal muscle of adult zebrafish are at comparable regenerative stages. (**A**) Heat maps showing expression patterns of *myog*, *myh7,* and *mymk* genes obtained by RNA-seq. (**B**) Relative expression levels of *myog*, *myh7,* and *mymk* were determined by RT-qPCR. Each bar represents the expression level of the indicated gene in stab-wounded or cryoinjured muscle relative to the corresponding non-injured control. NI, non-injured; stab, stab-wounded; CI, cryoinjured. n=4 for each experimental group. * P≤0.01. Mean Ct values ± SD are given in Supplementary Table S7.

Comparative analysis of DEGs revealed substantial overlap between the two injury types: 612 of 717 genes affected by stab wound (85%) were also differentially expressed in cryoinjured muscle (Fig. 6A; Supplementary Table S8). GO enrichment analysis showed that upregulated genes from both injury types (593 genes) shared a significant number of GO terms (88 biological process terms, 11 molecular function terms, 11 cellular component terms), mostly involved in ECM organization, immune response, regulation of cell differentiation, developmental growth, vasculature development, and others (Fig. 6B, Supplementary Table S9). Although the exact enriched GO terms differed between downregulated genes of the two datasets, both were associated with metabolic functions, particularly oxidoreductase activity, transmembrane transporter activity, phosphatase activity, and cofactor binding, indicating shared alterations in cellular energy metabolism in both injury types (Fig. 6B, Supplementary Table S9). Additionally, downregulated genes in cryoinjury were significantly enriched in terms associated with mature/functional muscle fiber (sarcomere, myofiber, contractile muscle fiber) (Fig. 6B, Supplementary Table S9).

**Fig. 6.**
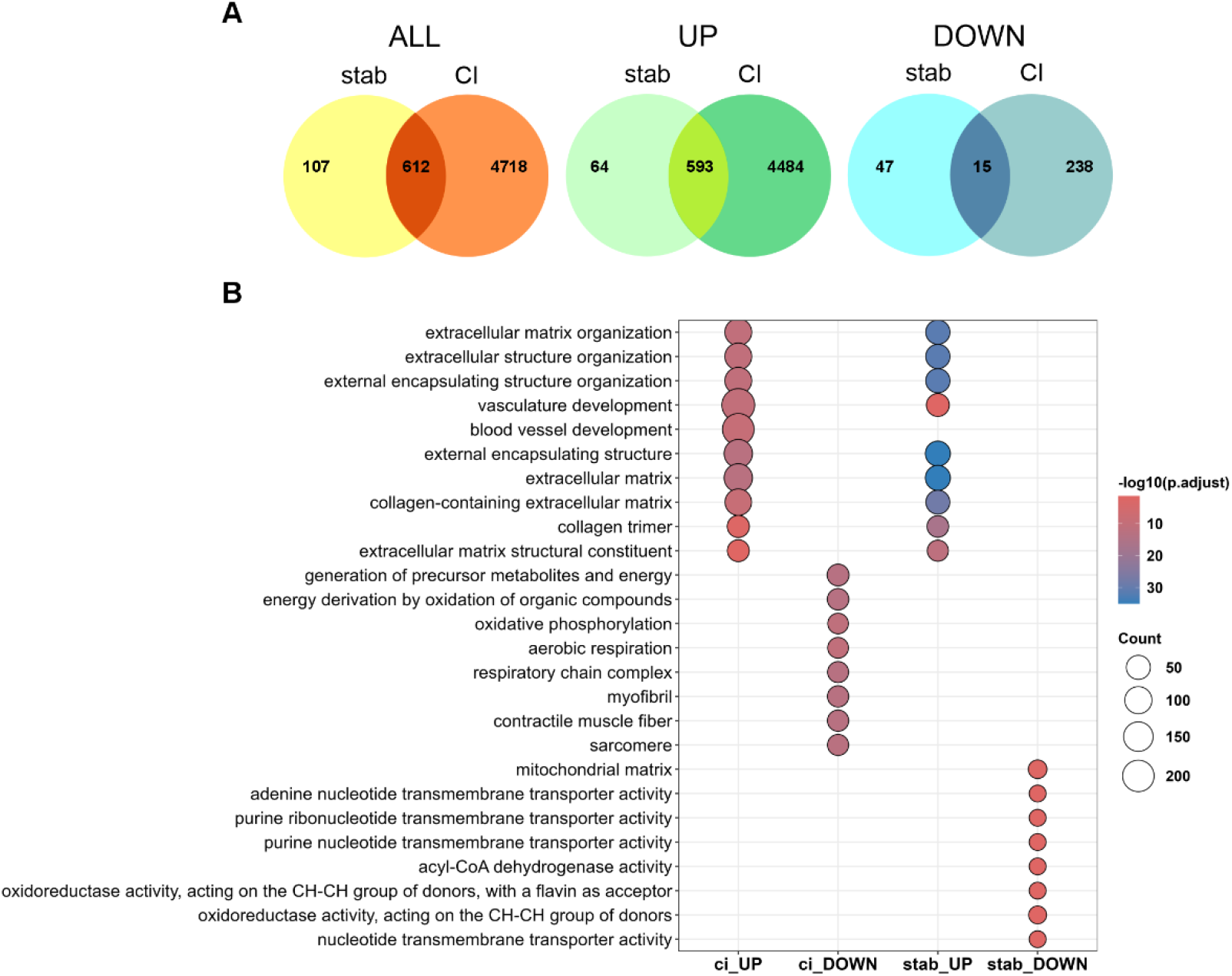
Shared DEGs and enriched GO terms in stab-wounded (stab) and cryoinjured (CI) skeletal muscle at 5 dpi and 7 dpci, respectively. (**A**) Venn diagram of all (ALL), upregulated (UP), and downregulated (DOWN) genes. (**B**) The dot plot of the top GO terms identified between up- and downregulated genes in stab-wounded and cryoinjured samples. The size of a dot represents the number of genes associated with the GO term, and the color of dots represents the -log10 of p.adjusted.

The principal KEGG pathways activated in both cryoinjured skeletal muscle at 7 dpci and stab-wounded muscle at 5 dpi included protein processing in the endoplasmic reticulum, focal adhesion, ECM-receptor interaction, phagosome, and apoptosis (Fig. 7A). Among the shared upregulated genes, Reactome pathway analysis revealed enrichment of terms related to ECM remodeling, including ECM organization, collagen formation, assembly of collagen fibrils, collagen biosynthesis, and ECM degradation, as well as immune-related processes, particularly MHC class II antigen presentation (Fig. 7B). In contrast, the 62 genes downregulated following stab injury showed no significant enrichment in either KEGG or Reactome pathways. By comparison, pathways downregulated in cryoinjured skeletal muscle were predominantly associated with cell motility and multiple energy metabolism processes, as described in the previous section (Fig. 3A).

**Fig. 7.**
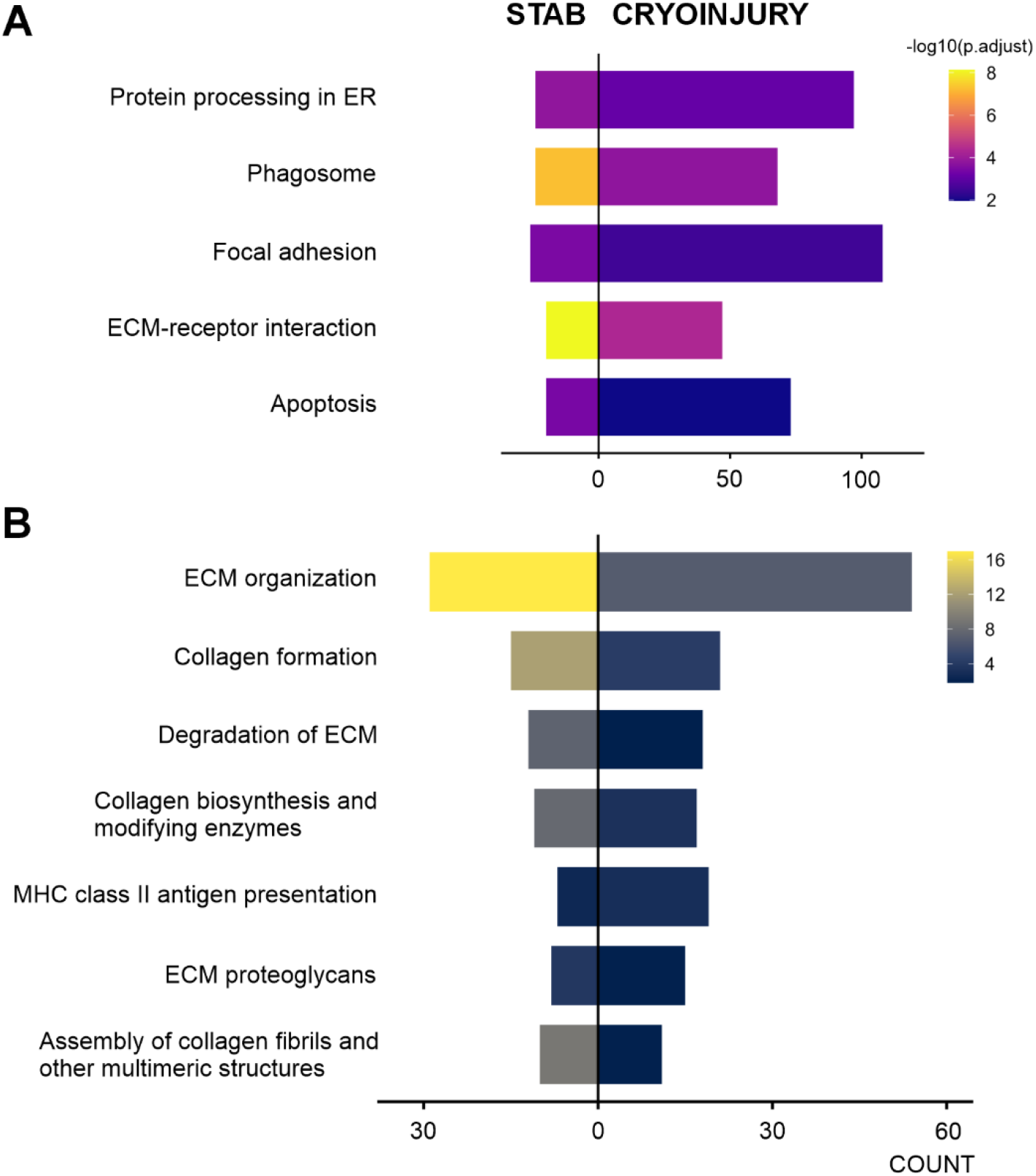
Bar plot of common KEGG (A) and Reactome (B) pathways enriched among DEGs upregulated following skeletal muscle injury induced by stab (left) and freezing (right).

PPI network analysis of upregulated DEGs shared between cryoinjured and stab-wounded skeletal muscle resulted in the identification of several functional clusters (Fig. 8, Supplementary Fig. S2, and Supplementary Table S10). Among the top five, Cluster 1 (score 11.17; 13 genes) was enriched for cell cycle-related terms, including DNA replication, mitotic spindle organization, and chromosome segregation, indicating robust activation of proliferative pathways. Two complementary clusters captured ECM dynamics, encompassing collagen biosynthesis and endoplasmic reticulum-mediated protein folding (Cluster 2, score 7.67; 13 genes), alongside integrin-mediated signaling and ECM organization (Cluster 3, score 4.67; 13 genes). A distinct immune-related cluster (Cluster 4, score 4.00; 4 genes) highlighted the regulation of myeloid leukocyte differentiation, particularly neutrophil lineage specification. Finally, a cluster enriched for myosin filament organization, motor proteins, and ER-associated protein processing (Cluster 5, score 3.47; 16 genes) indicated activation of muscle-specific differentiation programs and reassembly of the contractile apparatus.

**Fig. 8.**
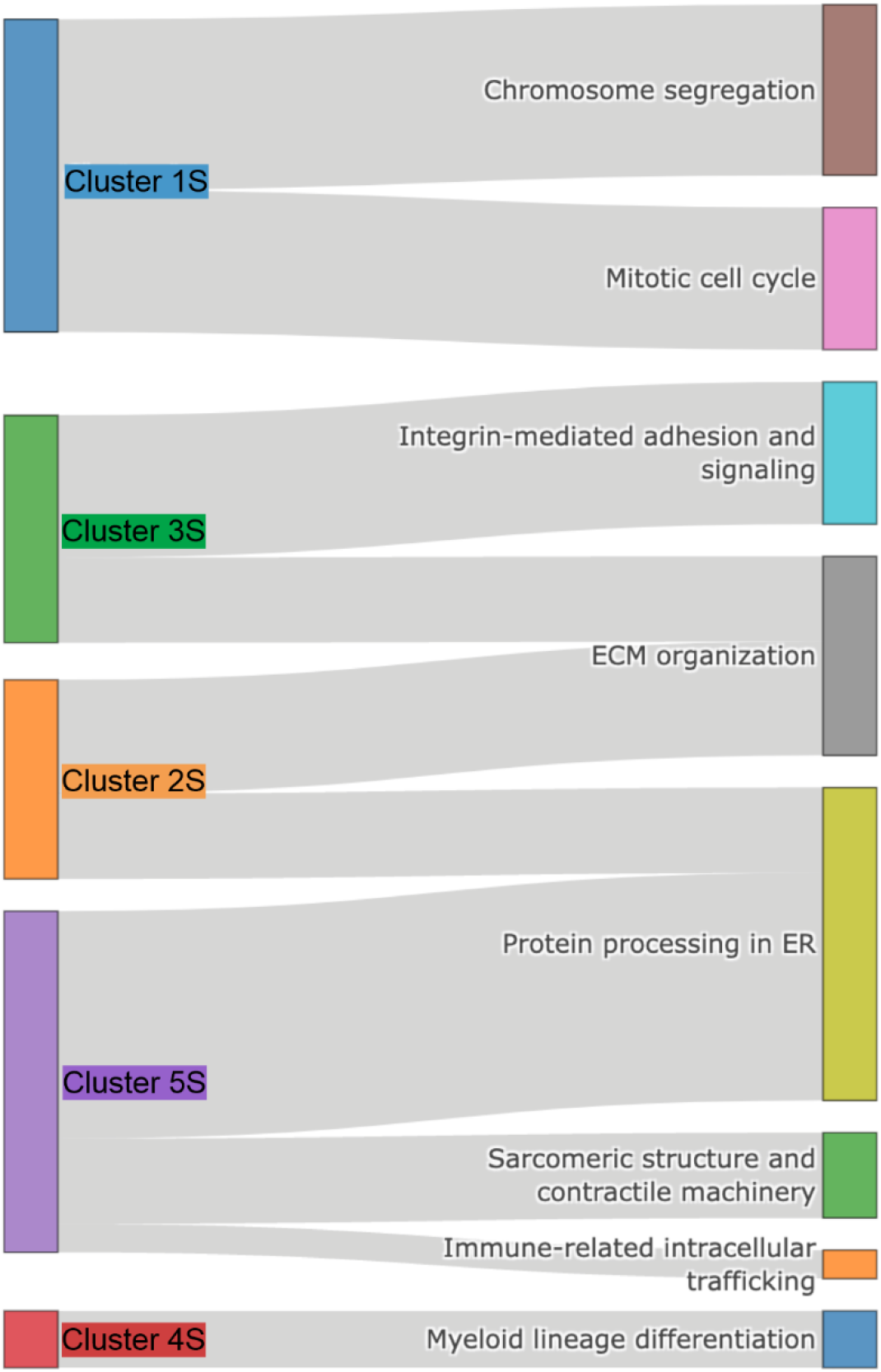
Sankey diagram showing connections between clusters from the PPI network analysis of DEGs shared between cryoinjured (at 7 dpci) and stab-wounded (at 5 dpi) adult zebrafish skeletal muscle and enrichment terms. Cluster 1S: Cell proliferation and mitotic machinery; Cluster 2S: ECM and protein processing; Cluster 3S: Cell-ECM interaction; Cluster 4S: Myeloid lineage differentiation and Cluster 5S: Muscle differentiation and ER proteostasis.

Collectively, these clusters delineate a shared, injury-independent regenerative framework characterized by tightly coordinated processes of cell proliferation, ECM production and signaling, immune regulation, and muscle fiber differentiation, underscoring the conserved nature of the skeletal muscle repair program in adult zebrafish.

## DISCUSSION

### Regenerative context of cryoinjured zebrafish skeletal muscle at 7 dpci

Cryoinjured skeletal muscle suffers from extensive loss of myofibers and disruption of tissue architecture. Zebrafish can repair this type of injury following a sequence of processes, including muscle degeneration, myogenic program activation, muscle restoration, and muscle remodeling (Oudhoff et al. 2024).

For comprehensive transcriptome profiling of cryoinjured zebrafish skeletal muscle, we selected a stage of repair at 7 dpci, which captures the transition from progenitor expansion toward myogenic differentiation and tissue rebuilding. It follows acute degeneration and inflammatory destruction, but precedes maturation and muscle remodeling. Along with nascent myofiber formation, this phase is also characterized by coordinated inflammation, immune response, ECM remodeling, and cell proliferation (Oudhoff et al. 2024). Our findings are in line with these reported results; at 7 dpci, immune-mediated debris clearance remains active, ECM and cell adhesion programs are highly dynamic, cell proliferation continues, and differentiation and fusion of myogenic precursors into immature fibers are in place. These observations are supported by upregulation of myogenic regulators, suppression of mature sarcomeric and metabolic genes, strong ECM, focal adhesion, and TGFβ signatures, enrichment of phagosome, lysosome, and inflammatory pathways, cell-cycle activation, actin and Rho GTPase-mediated remodeling, and reduced oxidative phosphorylation, indicating active reconstruction of skeletal muscle tissue, far from homeostasis.

Infiltration of nascent connective tissue at the injury site by newly forming muscle fibers (Oudhoff et al. 2024) is supported by our transcriptomics analysis. Ongoing myogenic differentiation is reflected in the upregulation of genes known to participate in the process of progenitor commitment and their regeneration-associated fusion, including master myogenic regulators myogenin (*myog*) and myogenic factor 5 (*myf5*), embryonic myosin isoform *myh7*, the cell-alignment mediators, junctional adhesion molecules (*jam*), and regulator of muscle fusion myomaker (*mymk*).

### Immune response in cryoinjured skeletal muscle

Persistent immune and phagocytic activity and inflammatory signaling in cryoinjured skeletal muscle at 7 dpci indicate ongoing tissue clearance and remodeling. Activated pathways are related to lysosomes, phagosomes, neutrophil degranulation, the innate immune system, antigen presentation, and platelet degranulation. This is consistent with histological findings of progressive accumulation of L-plastin-positive phagocytes during the first 10 days after injury, concomitant with tissue clearance and initiation of regeneration (Oudhoff et al. 2024). The enrichment of immune and phagocytic pathways observed in our transcriptomic dataset is also consistent with an active role of immune cells in tissue reconstruction beyond debris clearance alone, since it was proposed that infiltrating immune cells contribute to the activation of muscle stem cells and the establishment of a permissive regenerative niche (Tidball 2017; Chazaud 2020).

Additionally, cryoinjury activated the expression of Toll-like receptors (TLRs), likely through the release of damage-associated molecular pattern (DAMP) molecules from dying myofibers, a mechanism similar to that observed in the cryoinjured zebrafish heart (Vilahur and Badimón 2014). Myeloid differentiation factor 88 (*myd88*), with a role in neutrophil and macrophage recruitment and restriction of fibrotic response in the regenerating heart (Goumenaki et al. 2024), was also upregulated. In addition to mediating innate immune activation following cryoinjury, *myd88* may also function as a regulator of regenerative signaling pathways, since recently *myd88* signaling was shown to regulate the phosphoinositide 3 kinase (PI3K)/AKT pathway in the injured endocardium (Goumenaki et al. 2024). We found PI3K complex and PI3K regulatory activity as enriched terms in cryoinjured zebrafish skeletal muscle. Moreover, *cxcl18b*, a prominent inflammatory marker and transcriptional target of *myd88* (Goumenaki et al. 2024), was also upregulated. The concurrent activation of TLR signaling, PI3K-associated processes, and *cxcl18b* expression suggests that mechanisms previously implicated in zebrafish heart regeneration may also operate in skeletal muscle, representing a conserved, cross-tissue regenerative immune module. Persistent immune activity at 7 dpci appears to contribute to the establishment of a regenerative niche that supports extensive progenitor cell activation and expansion required for tissue reconstruction (Oudhoff et al. 2024).

### Cell expansion in cryoinjured skeletal muscle

Transcriptomic analysis identified a prominent proliferative module associated with DNA replication, mitotic cell-cycle progression, spindle assembly, chromosome segregation, and microtubule organization. Upregulation of cell-cycle regulators, including *pcna*, *mki67*, *foxm1*, *plk1*, *ccnb1*, *cdc20*, *bub1*, together with members of the minichromosome maintenance (MCM) complex, indicates that cell proliferation remains highly active at 7 dpci.

Among the most prominent members of this proliferative signature are cell-cycle regulators *foxm1* and *plk1*, whose orthologues have been implicated in mammalian skeletal muscle regeneration (Jia et al. 2019; Chen et al. 2020). Upregulation of *fosab*, *fosl1a*, and *fosl2* genes, encoding components of the AP-1 transcription factor complex, suggests rapid activation of an early injury-responsive transcriptional program. Together with increased expression of jun family members (*junbb*, *jund*), these genes likely contribute to coordinating multiple regenerative processes beyond cell-cycle regulation, including myogenic cell activation, ECM remodeling, inflammatory signaling, and tissue reconstruction (Almada et al. 2021; Endo 2023).

Although bulk RNA-seq does not resolve the identity of proliferating cell populations, our findings indicate that the regenerative environment at 7 dpci remains highly proliferative, with coordinated activation of cell-cycle regulators and transcriptional programs associated with myogenic progenitor function. This stage of repair appears to balance ongoing progenitor expansion with the initiation of myogenic differentiation, ensuring a sufficient cellular pool for efficient reconstruction of damaged muscle tissue. The conservation of these proliferative and myogenic regulatory programs between zebrafish and mammals further supports progenitor expansion as a fundamental component of vertebrate skeletal muscle repair.

### Myogenic differentiation in cryoinjured skeletal muscle

Upregulation of early myogenic regulators and fusion-associated genes in cryoinjured zebrafish skeletal muscle indicates active commitment of progenitor cells and formation of nascent myofibers. Muscle progenitor activation and recruitment to the injury site are evident from increased expression of the myogenic regulatory factor myogenin, which promotes expression of fusion-associated genes, including *mymk* (Ganassi et al. 2018). Myomaker and myomixer are conserved regulators of myoblast fusion in mammals and zebrafish (Millay et al. 2013; Landemaine et al. 2014; Zhang and Roy 2017; Shi et al. 2017; Bi et al. 2018). In contrast to myomaker, myomixer was not significantly induced, suggesting that regenerating tissue may not yet have reached peak multinucleated myofiber formation. Alternatively, fusion-associated genes may exhibit distinct temporal dynamics, as variability in myomaker and myomixer expression profiles has been reported among teleost regeneration models (Perelló-Amorós et al. 2022). Cryoinjury also induced expression of *jam2a* and *jam2b*, adhesion molecules involved in membrane protrusion formation during embryonic myoblast fusion in zebrafish (Luo et al. 2022), whereas *jam3b* remained unchanged. It appears that adult zebrafish skeletal muscle regeneration may not fully recapitulate the embryonic fusion program, although temporal differences in gene activation cannot be excluded.

Upregulation of embryonic *myh7* mirrors the appearance of nascent myofibers reported at 7 dpci (Oudhoff et al. 2024). Concurrent downregulation of several genes specific for mature muscle fibers further indicates that newly forming myofibers remain structurally immature and have not yet acquired a fully differentiated contractile phenotype. This transient suspension of mature muscle identity likely facilitates ongoing differentiation, fusion, and tissue reconstruction.

To conclude, cryoinjury activates a robust myogenic program that involves progenitor recruitment, differentiation, and the acquisition of fusion competence. However, the coexistence of developmental and adult muscle gene signatures suggests that newly formed fibers remain immature and are undergoing active structural remodeling. The following maturation of nascent myofibers occurs within a rapidly changing extracellular environment, making coordinated remodeling of the ECM an essential component of successful tissue reconstruction.

### ECM remodeling and mechanical reconfiguration in cryoinjured skeletal muscle

Cryoinjury of zebrafish skeletal muscle caused upregulation of genes coding for ECM constituents (collagens, laminins, fibronectin, fibromodulin, dermatopontin), activation of ECM remodeling factors (matrix metalloproteinases (MMPs) and tenascin C), and β-catenin, as an effector of Wnt signaling, known to be involved in the transition of skeletal muscle progenitor cells from proliferation to differentiation (Tanaka et al. 2011). Several induced ECM components, including fibromodulin and dermatopontin, have been linked to the regulation of myogenesis and promotion of skeletal muscle regeneration (Kim et al. 2019). Our results are in line with published findings that address wound clearance and collagen deposition in this exact cryoinjury model (Oudhoff et al. 2024). Consistent with the histological observations (Oudhoff et al. 2024), recruitment of neutrophils and macrophages likely promotes degradation of damaged matrix through cytokine and MMPs release, whereas activated muscle progenitors contribute to collagen synthesis required for restoration of tissue integrity (Gillies and Lieber 2011).

Extensive ECM remodeling is characterized by simultaneous matrix degradation and reconstruction. Beyond providing structural support for newly forming myofibers, this dynamic environment facilitates intercellular communication and coordination of regeneration (Ahmad et al. 2023). Importantly, it also generates an unstable mechanical landscape that drives biomechanical signaling and cytoskeletal reorganization (Calve and Simon 2012).

### Mechanotransduction in cryoinjured skeletal muscle

Skeletal muscle repair occurs within a dynamic mechanical environment, in which newly formed myofibers continuously adapt to forces generated by the surrounding intact muscle (Li et al. 2017). Following cryoinjury, extensive myofiber necrosis and ECM remodeling profoundly alter local mechanical tension (Zhou et al. 2020). Our results indicate that regenerating zebrafish skeletal muscle responds actively to these biomechanical cues through the coordinated activation of mechanotransduction pathways.

Upregulation of *piezo1* and *piezo2a.2* identifies Piezo channels as potential mechanosensitive gateways during muscle regeneration, suggesting that regenerating tissue experiences persistent mechanical stress associated with structural immaturity (Cox et al. 2017; Hirano et al. 2022). Additional induction of mechanosensitive *tmem63* (Kang and Lee 2024) and *trp* (Choi et al. 2020; Rolland et al. 2023) family members, together with the voltage-gated calcium channel *cav1* (Wang et al. 2024), indicates expansion and amplification of calcium-dependent mechanosensing. Increased expression of *mcur1* further suggests coupling of mechanically induced calcium influx with mitochondrial metabolism, linking mechanical stress detection to the energetic demands of tissue reconstruction (Vais et al. 2015; Haschke et al. 2025). Consistent with this, regenerating skeletal muscle exhibited strong activation of the RhoA/ROCK pathway (Nishiyama et al. 2004; Brondolin et al. 2023), together with extensive remodeling of integrin-mediated adhesion complexes through induction of integrins, talins, kindlins, and calpains. Because Piezo-mediated calcium influx activates calpains that regulate integrin–cytoskeleton coupling (Göll et al. 2003; Franco et al. 2004; Coste et al. 2010), these findings further point to extensive cytoskeletal reorganization and adhesion remodeling in response to altered mechanical demands.

Upregulation of several Hippo-YAP pathway components, including *yap1*, *stk3*, *rhoa*, *rock*, and the mechanosensitive target *ankrd1a*, highlights coordinated regulation of nuclear mechanotransduction during regeneration (Dasgupta and McCollum 2019; Driskill and Pan 2023; Guo et al. 2024). Particularly notable is the induction of *ankrd1a*, a YAP target and titin-associated mechanosensor (Miller et al. 2003; Dupont et al. 2011). Combined with our previous finding that *ankrd1a* mutants exhibit accelerated stab wound repair (Milovanovic et al. 2025), these results suggest that the YAP–ANKRD1A axis can modulate regenerative progression and may function as a mechanical checkpoint coordinating differentiation, fusion, and maturation of newly formed myofibers.

Finally, enrichment of PI3K–mTOR signaling terms indicates increased anabolic activity accompanying tissue reconstruction. As a central regulator of mechanically induced protein synthesis (Hornberger 2011; Wei et al. 2019), mTOR signaling likely supports the production of structural components required for rebuilding regenerating myofibers. This interpretation is supported by previous evidence that pharmacological inhibition of mTOR impairs regeneration of cryoinjured zebrafish skeletal muscle (Oudhoff et al. 2024). Together, these findings highlight the close integration of mechanotransduction with anabolic signaling, linking mechanical adaptation to the extensive biosynthetic reprogramming required for ongoing tissue reconstruction.

### Cellular logistics during skeletal muscle repair

Successful skeletal muscle repair requires coordinated synthesis, quality control, and intracellular distribution of newly produced proteins to support myogenic differentiation, myofiber growth, and tissue reconstruction (Chargé and Rudnicki 2004).

Biosynthetic expansion represents the production arm of skeletal muscle repair. Consistent with increased anabolic demands, cryoinjured skeletal muscle exhibited strong activation of ribosome biogenesis, transcriptional and translational machinery, including induction of the *myc* regulatory network components, ribosomal proteins, RNA polymerase subunits, nucleolar ribosome-biogenesis factors, translation initiation factors, *rsk* family members, and *mtor* (Cho et al. 2007; Hornberger 2011; Roméo et al. 2011; Xu and Liu 2026; Cui et al. 2026). These findings indicate extensive expansion of biosynthetic capacity and agree with recent evidence that *myc* activation and ribosome biogenesis are conserved features of skeletal muscle regeneration in mice and humans (Cui et al. 2026).

Proteostasis constitutes the quality-control arm of skeletal muscle repair. Alongside increased biosynthesis, regenerating muscle activated autophagy, lysosomal pathways, polyamine metabolism, ER stress responses, and protein-folding machinery, by upregulation of *gabarap*, *atg* family members, *sirt1*, *xbp1*, *p4hb*, *calr*, and *dnaJ/hsp40* genes (Mizushima et al. 2011; Minois et al. 2012; Tang and Rando 2014). Together with previous evidence of p62 accumulation during wound clearance (Oudhoff et al. 2024), these findings indicate coordinated protein turnover, organelle recycling, and proteome quality control that support proliferative expansion, myogenic differentiation, and tissue remodeling (Hetz and Papa 2017; Ceci et al. 2022).

Intracellular trafficking functions as the deployment arm of skeletal muscle repair. We identified enrichment of terms related to Golgi-to-ER trafficking, coated vesicles, clathrin-mediated transport, nucleocytoplasmic transport, and cytoskeletal remodeling, together with increased expression of genes mediating vesicular transport and membrane dynamics (*copa, copb1, gosr2, bet1l, rab8a, snx2, snapin, cdc42, actr3, arpc3, wasf2, wipf1b*). These systems likely coordinate the delivery of newly synthesized proteins, ECM components (Bonnans et al. 2014; Malhotra and Erlmann 2015), and membrane proteins required for myoblast fusion (Kim et al. 2015), mechanotransduction and adhesion (Schwartz and DeSimone 2008; Demonbreun and McNally 2017).

Collectively, these findings suggest that successful zebrafish skeletal muscle regeneration relies on a coordinated cellular logistics network in which biosynthetic expansion, proteostasis, and intracellular trafficking function as complementary arms supporting efficient tissue reconstruction.

### Integrated conceptual model of skeletal muscle repair at 7 dpci

One of the most striking findings of the present study is the extensive temporal overlap of regenerative processes at 7 dpci in zebrafish skeletal muscle. Skeletal muscle regeneration is generally described as a sequence of distinct phases or, more recently, as a ”wave-on-wave” process (Forcina et al. 2020); however, our transcriptomic data reveal the simultaneous activation of immune remodeling, cell expansion, myogenic differentiation, ECM reconstruction, mechanotransduction, biosynthetic adaptation, proteostasis, and intracellular trafficking. These findings suggest that zebrafish skeletal muscle regeneration is a highly coordinated systems-level process in which multiple biological modules operate in parallel. At 7 dpci, cryoinjured muscle remains far from homeostasis and functions as an integrated regenerative system, where immune-mediated tissue clearance, cell expansion, tissue rebuilding, mechanical adaptation, and biosynthetic reprogramming are tightly interconnected and collectively drive tissue reconstruction (Fig. 9).

**Fig. 9.**
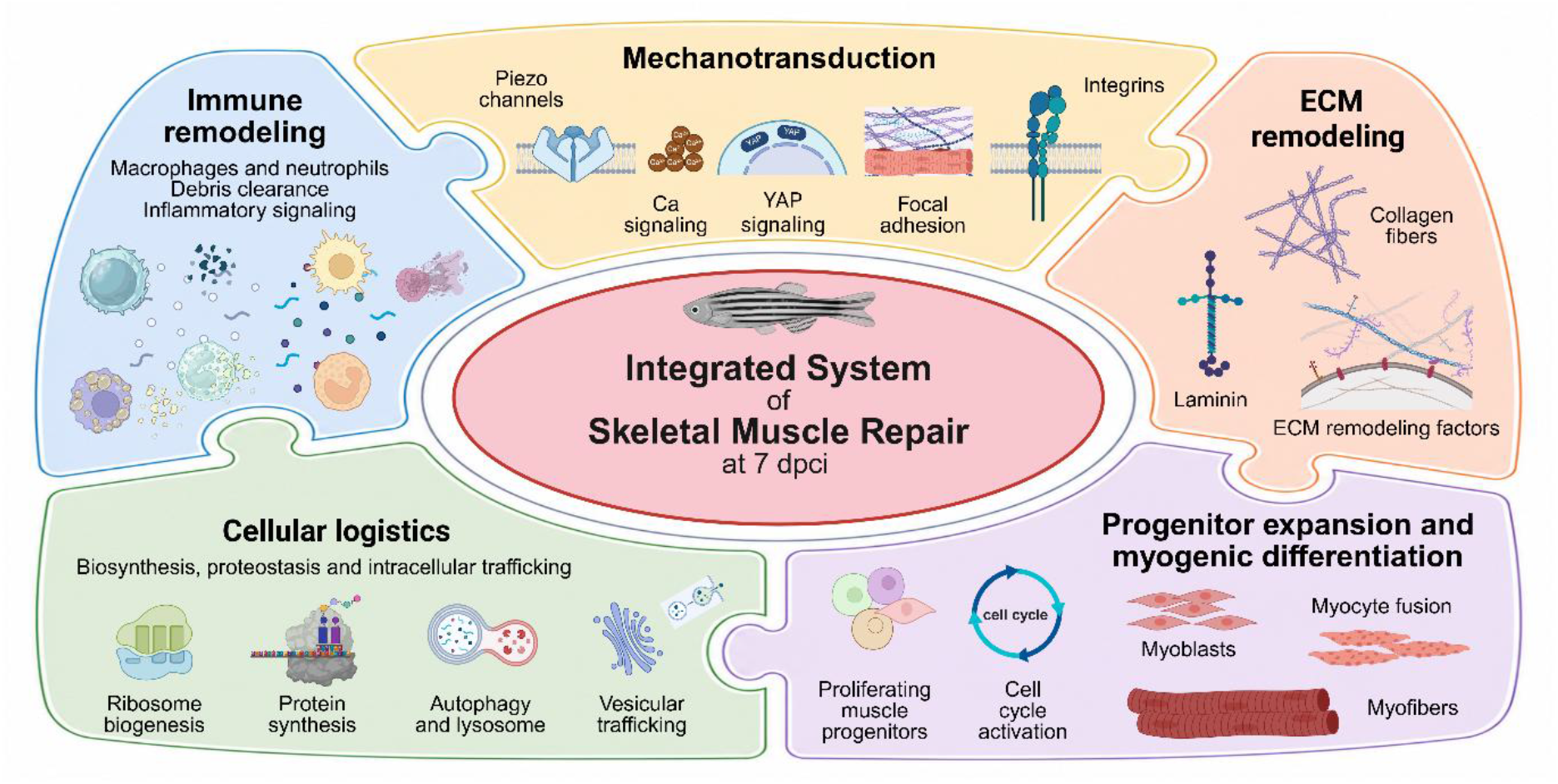
Integrated conceptual model of cryoinjured zebrafish skeletal muscle at 7 dpci. Cryoinjured zebrafish skeletal muscle functions as an integrated regenerative system in which immune remodeling, progenitor expansion, myogenic differentiation, ECM reconstruction, mechanotransduction, and cellular logistics operate simultaneously and interact extensively to drive tissue reconstruction. Together, these interconnected biological modules form a conserved regenerative architecture that underlies successful skeletal muscle repair. Created in BioRender. Milošević, E. (2026) https://BioRender.com/l3yv2n8.

Within this framework, the individual biological modules do not operate as independent pathways but as highly interconnected components of a coordinated regenerative network. Immune remodeling establishes a permissive regenerative niche that supports progenitor activation, while proliferative and differentiation programs drive continuous generation of new myogenic cells and formation of nascent myofibers. ECM remodeling reconstructs the structural and mechanical environment, mechanotransduction integrates these evolving biomechanical cues with cellular behavior, and biosynthetic expansion, proteostasis, and intracellular trafficking function together as a cellular logistics network that supplies, quality controls, and distributes the molecular components required for tissue reconstruction.

Collectively, these observations support an integrated conceptual model in which successful skeletal muscle repair emerges from dynamic interactions among specialized but tightly interconnected biological modules. This systems-level organization provides a framework for understanding how skeletal muscle progressively restores its structure, function, and homeostasis following extensive tissue damage.

### A shared transcriptional response in cryoinjured and stab-wounded skeletal muscle reveals a conserved regenerative program

Comparison of phase-matched regenerating skeletal muscle following cryoinjury and stab injury revealed a striking degree of transcriptional overlap. Although cryoinjury induced a substantially larger number of DEGs than stab injury, the majority of genes affected by stab injury were also differentially expressed following cryoinjury. This observation indicates that distinct injury modalities converge on a common regenerative program despite major differences in the extent and nature of tissue damage.

Importantly, the shared response was not limited to individual genes but encompassed coherent biological modules. PPI network analysis of overlapping upregulated genes identified clusters associated with cell proliferation, ECM production and signaling, immune regulation, muscle differentiation, and endoplasmic reticulum–associated protein processing. These modules closely correspond to the major biological processes identified throughout the cryoinjury dataset, supporting the existence of a conserved regenerative program that is activated independently of injury modality. Future studies should answer the question of how the zebrafish regulatory landscape successfully decouples these pathways from the fibrotic outcomes observed in mammals (Gallardo et al. 2025).

The asymmetry between the two injury models is particularly informative. While stab injury activated a relatively limited transcriptional response, cryoinjury elicited a much broader transcriptomic reorganization. Nevertheless, the shared genes and pathways indicate that severe injury does not trigger a fundamentally distinct regenerative mechanism. Rather, cryoinjury appears to recruit the same core regenerative architecture together with a substantial expansion of transcriptional programs required to accommodate extensive tissue destruction, necrotic tissue clearance, ECM reconstruction, and large-scale tissue rebuilding. Thus, injury severity appears to influence the breadth of transcriptional engagement rather than the fundamental identity of the regenerative response.

The extensive overlap was primarily observed among upregulated genes, whereas downregulated genes showed considerably less conservation. This suggests that the activation of regenerative processes is highly stereotyped, whereas the suppression of physiological tissue functions depends more strongly on the severity of injury and tissue damage. Consistent with this interpretation, larger injuries in zebrafish recruit more Pax7^+^ progenitors than focal lesions (Knappe et al. 2015), indicating that the broader activation of myogenic, inflammatory, ECM, and mechanotransduction pathways observed after cryoinjury likely reflects greater regenerative demand rather than activation of a distinct regenerative mechanism.

Collectively, these findings support a model in which adult zebrafish skeletal muscle regeneration is governed by a conserved regenerative architecture that can be deployed across diverse injury contexts. Stab injury appears to activate a more restricted regenerative program sufficient for efficient tissue repair, whereas cryoinjury engages a quantitatively expanded version of the same regenerative machinery.

Similar conclusions were reached regarding zebrafish heart regeneration, where a core regenerative program shared across three injury models was identified, together with injury-specific responses (Botos et al. 2023). The close parallel with our skeletal muscle data suggests that regeneration of both zebrafish skeletal muscle and heart is governed by a conserved regenerative architecture upon which injury-specific adaptations are superimposed according to regenerative demand. The preservation of these core regenerative modules highlights the intrinsic robustness of zebrafish regeneration despite substantial differences in injury modality.

### Perspectives

The present study extends the molecular characterization of adult zebrafish skeletal muscle repair after cryoinjury by providing the first comprehensive transcriptomic analysis of this injury model and the first comparison of regenerative programs across two distinct injury modalities. Together, these findings expand the utility of the adult zebrafish skeletal muscle model for systems-level investigation of successful regeneration.

In contrast to mammalian skeletal muscle, where severe injury is frequently associated with persistent fibrosis, incomplete regeneration, and functional impairment, zebrafish efficiently progress through regenerative stages while transiently reactivating developmental-like programs associated with tissue growth and muscle formation (Pfefferli and Jaźwińska 2017; Oudhoff et al. 2024). Characterization of the molecular networks underlying this regenerative competence may therefore help identify conserved mechanisms that support efficient tissue restoration and provide insights into pathways that are limited or dysregulated in mammalian muscle repair.

At the same time, the molecular programs identified here are likely distributed across multiple cellular populations and dynamically regulated throughout the repair process. While the present work provides a detailed transcriptomic snapshot of a critical regenerative stage, an important unresolved question is whether the extensive overlap of immune, myogenic, ECM, mechanotransductive, and biosynthetic programs observed at 7 dpci represents a unique state of maximal regenerative integration or a more persistent feature of tissue repair. Defining how these biological modules are reorganized, resolved, or maintained throughout regeneration will be essential for understanding not only the temporal architecture of skeletal muscle repair, but also the general principles that enable successful vertebrate tissue regeneration. Future studies spanning multiple phases of repair and integrating single-cell, spatial transcriptomic, and proteomic approaches will be necessary to resolve the cellular origin, spatial organization, and functional interactions underlying successful tissue reconstruction. Combined with the unique experimental advantages of the zebrafish model, these approaches may help define fundamental principles of vertebrate tissue regeneration and ultimately inform regenerative strategies for mammalian skeletal muscle repair.

## SUPPLEMENTARY INFORMATION

**Supplementary Fig. S1**. PPI network modules identified among downregulated DEGs after cryoinjury (https://doi.org/10.6084/m9.figshare.33197403).

**Supplementary Fig. S2.** PPI network modules identified among upregulated DEGs shared between cryoinjured (at 7 dpci) and stab-wounded (at 5 dpi) adult zebrafish skeletal muscle (https://doi.org/10.6084/m9.figshare.33197427).

**Supplementary Table S1**. List of primers used for RT-qPCR (https://doi.org/10.6084/m9.figshare.33196863).

**Supplementary Table S2**. List of DEGs in cryoinjured zebrafish skeletal muscle at 7 dpci (https://doi.org/10.6084/m9.figshare.33196914).

**Supplementary Table S3**. List of GO terms enriched among upregulated and downregulated DEGs in cryoinjured zebrafish skeletal muscle at 7 dpci (https://doi.org/10.6084/m9.figshare.33196938).

**Supplementary Table S4**. List of KEGG pathways enriched among upregulated and downregulated DEGs in cryoinjured zebrafish skeletal muscle at 7 dpci (https://doi.org/10.6084/m9.figshare.33196953).

**Supplementary Table S5**. List of Reactome pathways enriched among upregulated and downregulated DEGs in cryoinjured zebrafish skeletal muscle at 7 dpci (https://doi.org/10.6084/m9.figshare.33196962).

**Supplementary Table S6**. PPI network analysis of upregulated and downregulated DEGs in cryoinjured zebrafish skeletal muscle at 7 dpci (https://doi.org/10.6084/m9.figshare.33196965).

**Supplementary Table S7**. Average Ct values ± SD for *myog*, *myh7*, *mymk,* and *rpl13a* transcript fragments amplified by RT-qPCR (https://doi.org/10.6084/m9.figshare.33197163).

**Supplementary Table S8**. List of shared DEGs between cryoinjured skeletal muscle at 7 dpci and stab-wounded skeletal muscle at 5 dpi (https://doi.org/10.6084/m9.figshare.33197199).

**Supplementary Table S9**. GO terms associated with upregulated and downregulated DEGs in cryoinjured (at 7 dpci) and stab-wounded (at 5 dpci) zebrafish skeletal muscle (https://doi.org/10.6084/m9.figshare.33197205).

**Supplementary Table S10**. PPI network analysis of upregulated DEGs shared between cryoinjured (at 7 dpci) and stab-wounded (at 5 dpci) zebrafish skeletal muscle (https://doi.org/10.6084/m9.figshare.33197220).

## DATA AVAILABILITY

The RNA-seq data reported and used in this manuscript have been deposited in the Gene Expression Omnibus (GEO) database with the accession numbers GSE319584 and GSE277480. Other relevant data can be found within the article and supplementary information.

